# A chromosome-scale hybrid genome assembly of the extinct Tasmanian tiger (*Thylacinus cynocephalus*)

**DOI:** 10.1101/2022.03.02.482690

**Authors:** Charles Feigin, Stephen Frankenberg, Andrew Pask

## Abstract

The extinct Tasmanian tiger or thylacine (*Thylacinus cynocephalus*) was a large marsupial carnivore native to Australia. Once ranging across parts of the mainland, the species remained only on the island of Tasmania by the time of European colonization. It was driven to extinction in the early 20^th^ century and is an emblem of native species loss in Australia. The thylacine was a striking example of convergent evolution with placental canids, with which it shared a similar skull morphology. Consequently, it has been the subject of extensive study. While the original thylacine assemblies published in 2018 enabled the first exploration of the species’ genome biology, further progress is hindered by the lack of high-quality genomic resources. Here, we present a new chromosome-scale hybrid genome assembly for the thylacine, which compares favorably with many recent *de novo* marsupial genomes. Additionally, we provide homology-based gene annotations, characterize the repeat content of the thylacine genome and show that, consistent with demographic decline, the species possessed a low rate of heterozygosity even compared to extant, threatened marsupials.

**Significance:** The lack of high-quality genomes for extinct species inhibits research into their biology. Moreover, marsupials are underrepresented among sequenced genomes. Here, we present a new, chromosome-scale thylacine genome. This high-quality assembly is a valuable new resource for studies on marsupial carnivores.

## Introduction

The Tasmanian tiger or thylacine (*Thylacinus cynocephalus*; Fig. 1a) was the largest marsupial predator of the Holocene (Mitchell, et al. 2014; Prowse, et al. 2014). While it once inhabited mainland Australia, by the arrival of European colonists it was restricted to the island of Tasmania (Lambeck and Chappell 2001; Paddle 2000). The thylacine was considered an agricultural pest and targeted by an extermination campaign, incentivized by a £1 bounty (Fig. 1b). The last known individual died in 1936 and the species was declared extinct in 1986 (Paddle 2000). The thylacine was captured in multiple photographs and short films, contributing to its status as an emblem of Australia’s high extinction rate among native species (Sleightholme and Campbell 2018; Woinarski, et al. 2015).

**Fig. 1.**
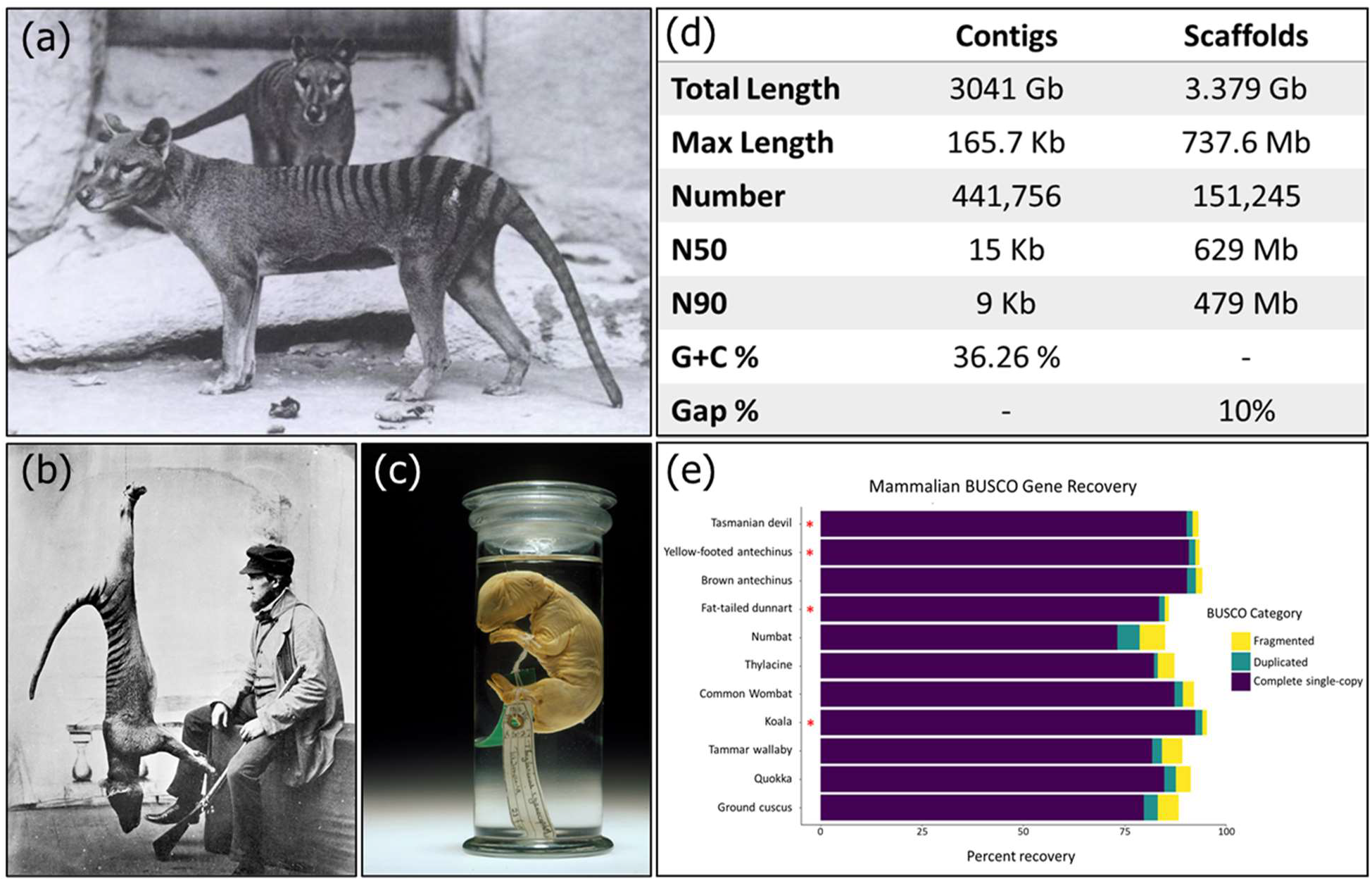
(a) Adult thylacines in captivity. The thylacine was noted for its canid-like morphology. (b) A wild thylacine killed by a hunter. A bounty on thylacines contributed to their extinction. (c) Thylacine pouch young specimen C5757 (Melbourne Museum; Victoria, Australia) provided DNA used for genome sequencing. (d) Assembly metrics for the improved thylacine genome. (e) Comparison of BUSCO gene recovery from the thylacine genome and several recently-released marsupial assemblies. Asterisk indicates assemblies incorporating long reads.

The relative abundance of thylacine specimens in museums has facilitated extensive study of its morphology, ecology and evolution (Newton, et al. 2018; Rovinsky, et al. 2021; White, et al. 2018; Wroe, et al. 2007). Recently, it has also become a focal species for genomic research, with the first genome assemblies being published in 2018, using DNA from a >100-year-old ethanol-preserved pouch young specimen (Fig. 1c) (Feigin, et al. 2018). These assemblies were used to explore the molecular basis of thylacine-canid craniofacial convergence, confirm its phylogenetic relationships, and infer its demographic history (Feigin, et al. 2018). Subsequent studies examined enhancer evolution and characterized the thylacine’s immune gene complement (Feigin, et al. 2019; Peel, et al. 2021). However, contiguity of the original assemblies was limited by the fragmentary nature of historical DNA and the absence of high-quality assemblies from related species suitable for reference-guided scaffolding (Feigin, et al. 2018). This presents a substantial challenge for continued research into the thylacine’s genome biology (Garrett Vieira, et al. 2020; Peel, et al. 2021).

The thylacine (family Thylacinidae) represents the closest sister lineage to the families Dasyuridae and Myrmecobiidae (Feigin, et al. 2018; Miller, et al. 2009; Mitchell, et al. 2014). These groups contain numerous species of significant interest to evolutionary, developmental and conservation biology, such as the Tasmanian devil, quolls, dunnarts and the numbat (Cook, et al. 2021; Fancourt 2016; Spencer, et al. 2020; Stahlke, et al. 2021; Wright, et al. 2020). Moreover, the thylacine’s exceptional craniofacial similarities with canids, despite their ~160 million year divergence, make the species an excellent model system to study the genomic basis of morphological evolution (Bininda-Emonds, et al. 2007; Feigin, et al. 2018; Newton, et al. 2021; Rovinsky, et al. 2021). Improved genomic resources for this species are thus of considerable value to the broader genomics community. Here, we leveraged improvements in short read assembly tools and newly-available marsupial reference genomes to produce a chromosome-scale hybrid genome assembly for the thylacine.

## Results and Discussion

### Genome Assembly and Assessment

The new thylacine assembly is composed of 7 large scaffolds, corresponding to each of the 6 dasyuromorph autosomes and the X chromosome (Supplementary Table 1), together comprising ~93.25% of the sequence content (Deakin 2018). The gap-free assembly size is ~3.04Gbp and G+C content is 36.26%, comparable to that of the Tasmanian devil (Fig. 1d, Supplementary Table 2). Scaffold N50 and N90 are high (629Mbp and 479Mbp respectively), reflecting the large size of dasyuromorph autosomes (Deakin 2018). Contig N50 was 5-fold higher than that of the original *de novo* draft assembly, and similar to that of several other recent marsupial assemblies (Supplementary Table 2). A tail of small scaffolds comprising approximately 205Mbp remained unplaced, contributing to a relatively high gap percentage (~10%; Fig. 1d). Nonetheless, the new assembly represents a dramatic improvement in contiguity.

To evaluate the completeness and integrity of the assembly, BUSCO was used to annotate benchmarking mammalian orthologs. This identified 82.3% of BUSCO genes as complete and single-copy, with little duplication (0.9%). Another 4.1% were found as partial copies (Fig. 1e). This is a drastic increase over the original thylacine *de novo* assembly, from which BUSCO recovery was negligible (<10%), owing to low contiguity (Supplementary Table 3). While BUSCO gene recovery compares well with several other recently released marsupial assemblies, particularly those built from short read-based contigs scaffolded with Hi-C, it lags somewhat behind a small number of assemblies built using long reads and Hi-C (Fig. 1e, Supplementary Table 4). Unfortunately, the century-long room-temperature preservation of all existing thylacine tissue samples, and corresponding DNA fragmentation, limits the potential for long read sequencing to be applied productively in this species.

### Repeat Classification and Genome Annotation

Repetitive regions in the thylacine genome were annotated with RepeatMasker, using a custom database of species-specific and curated marsupial repeats (Fig. 2a) (Ellinghaus, et al. 2008; Flynn, et al. 2020; Hubley, et al. 2016; Tarailo-Graovac and Chen 2009). Interspersed repeats constituted ~56% of the assembly (Supplementary Table 5). Consistent with the highly conserved genome organization of dasyuromorphs, the thylacine had similar overall repeat composition to its living relatives (Tian, et al. 2022). The dominant repeat class was LINE elements (~36.5%), occurring at a frequency comparable to that of the Tasmanian devil (~39%), though somewhat lower than that of the brown antechinus (~45%) (Tian, et al. 2022).

**Fig. 2.**
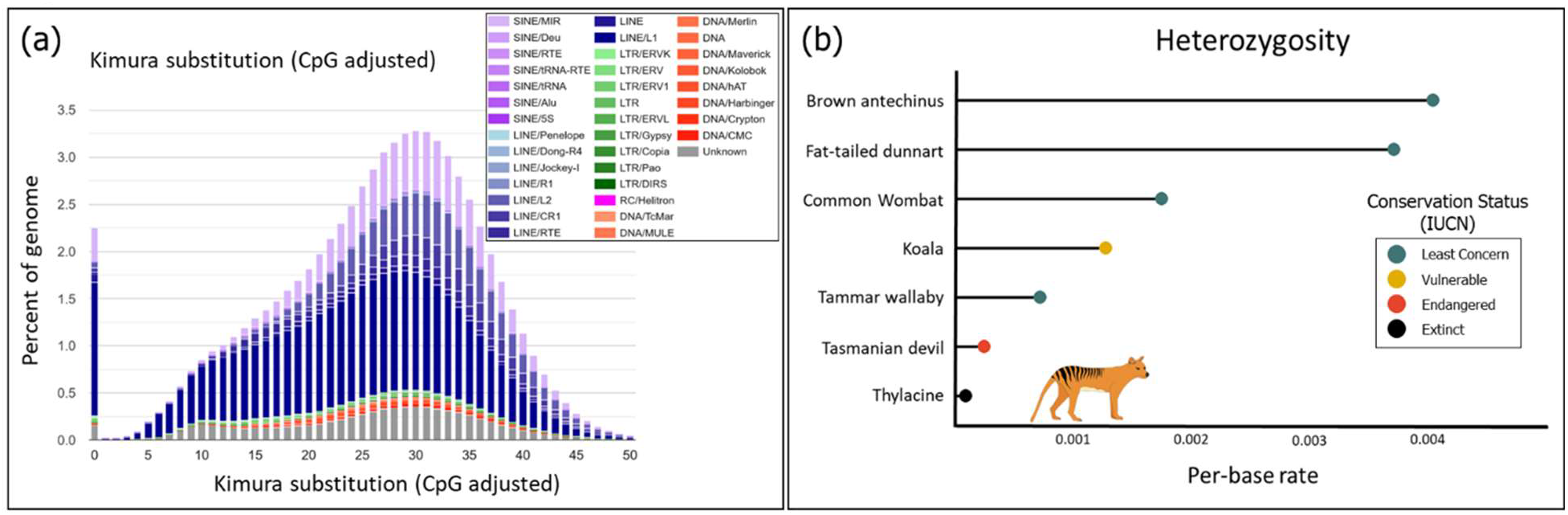
(a) Interspersed repeat landscape of thylacine genome. The percentage of total genome size and sequence divergence (based on CpG-adjusted Kimura substitution level) are shown for each repeat subclass. (b) Comparison of the per-base rate of heterozygosity in the thylacine and several extant marsupials. The thylacine showed the lowest heterozygosity of examined marsupial species.

Interestingly, we observed that LTRs were sparse in the thylacine genome (~1.51%) compared to previously studied marsupial species (which ranged from 6.53%-18.89%; Supplementary Table 5) (Tian, et al. 2022).

To provide gene annotations for the new thylacine assembly, we identified orthologs to Tasmanian devil genes using a homology-based annotation liftover procedure (see Methods and Methods). Ortholog recovery was high, with ~96% of gene models being successfully transferred to the thylacine genome, comparable to or exceeding that of other dasyuromorphs (Supplementary Table 6). Interestingly, we observed disparities in the detection of different short RNA classes. In particular, micro-RNAs (miRNAs) showed nearly complete recovery from the thylacine genome (~98%), compared with ~71% of small nucleolar RNAs and just ~37% of small nuclear RNAs (snoRNAs and snRNAs respectively; Supplementary Table 6). A similar pattern was observed among other dasyuromorphs, which showed lower snoRNA and snRNA recovery (particularly in species more distantly-related to the Tasmanian devil), while generally retaining high miRNA recovery (Supplementary Table 6). Taken together, this suggests that while many miRNAs are ancestral to Dasyuromorphia (hence having orthologs across species) and have remained conserved over time, the evolution of snRNAs and snoRNAs in this lineage has potentially been more dynamic, with accelerated sequence divergence and/or more rapid turnover of individual elements among species.

### Genetic Diversity

We next sought to gain insights into the thylacine’s genetic diversity prior to its extinction. Previously, multiple sequentially Markovian coalescent (MSMC) analysis was used to infer the demographic history of the thylacine. This uncovered evidence of an extended period of genetic decline predating the arrival of humans in Australia and the thylacine’s isolation on Tasmania (Feigin, et al. 2018; Schiffels and Durbin 2014). A decrease in genetic diversity concomitant with such demographic decline may have left the thylacine vulnerable to inbreeding depression, reducing its fitness on the backdrop of pressures imposed by humans. To further explore this possibility, heterozygosity was calculated in non-repetitive regions of the thylacine genome and compared to that of extant marsupials with varying conservation statuses. Consistent with reduced genetic diversity preceding its extinction, the thylacine had the lowest rate of heterozygosity among the marsupials examined, including vulnerable or endangered species (Fig. 2b, Supplementary Table 7).

## Conclusions

The quality of the first draft thylacine assemblies limited their utility in genomic research. Gene recovery was severely impaired by low contiguity, and repetitive regions were not adequately represented (Feigin, et al. 2018). By contrast, our new thylacine genome has a ~5-fold larger contig N50, comparable to that of many recent marsupial assemblies. Moreover, we have produced chromosome-scale scaffolds that enable the recovery of numerous genetic elements with orthologs in related species. This assembly has also permitted the first examination of the repeat composition and heterozygosity of the thylacine genome. Future whole-genome resequencing studies, empowered by this assembly, have the potential to provide population-level insights into the thylacine’s demography and level of genetic load prior to its extinction.

## Materials and Methods

### Genome Assembly

Thylacine reads were accessed from NCBI Sequence Read Archive (SRA; Supplementary Table 8). These data originated from individual C5757, which we previously used to produce the original contig-level *de novo* assembly and a read-mapping-based, reference-guided assembly of non-repetitive regions (Feigin, et al. 2018).

*De novo* contigs were assembled using MEGAHIT v1.2.9 (Li, et al. 2015) with multiple k-mer lengths (kmers = 21, 29, 39, 59, 79, 99, 119, 141). Purging of redundant haplotypes and short read scaffolding were performed using Redundans v0.14a (parameters: identity = 0.8, overlap = 0.8, minLength = 200bp, joins = 5, limit = 1.0, iterations = 2) (Pryszcz and Gabaldón 2016). Purging removed ~178.5Mbp of sequence.

Dasyuromorphs possess an exceptionally-conserved karyotype (2n = 14), with nearly identical chromosome sizes and g-banding patterns (Deakin 2018; Rofe and Hayman 1985). Moreover, sequence mapability between thylacine and Tasmanian devil is high (Feigin, et al. 2018). Therefore, chromosome-scale thylacine scaffolds were produced by ordering thylacine *de novo* scaffolds and inferring gap sizes through alignment against the recently-available Tasmanian devil reference genome (GCF_902635505.1/mSarHar1.11; (O’Leary, et al. 2016)) using RagTag v2.1.0 (RagTag parameters: scaffold, -f 200, -r, -g 100 -m 10000000; minimap2 v2.22-r1101 parameters: -x asm 10) (Alonge, et al. 2021; Alonge, et al. 2019; Li 2018).

### Genome Annotation

Repeat elements were annotated using RepeatMasker v4.1.2 (Flynn, et al. 2020; Tarailo-Graovac and Chen 2009). Custom thylacine repeat libraries were produced with RepeatModeler v2.0.2a and LTRharvest v1.6.2, and were combined with marsupial repeats contained with the Dfam3.2 database (Ellinghaus, et al. 2008; Flynn, et al. 2020; Hubley, et al. 2016). RepeatMasker was then run on each chromosome using this library (Supplementary Table 5). The repeat landscape of the thylacine genome was visualized using the calcDivergenceFromAlign.pl and createRepeatLandscape.pl scripts provided with RepeatMasker. This displays the genome percentage of each repeat subclass, organized by CpG-adjusted kimura substitution level (a distance-based proxy for repeat copy age) (Flynn, et al. 2020; Kimura 1980).

Given the thylacine’s extinction, RNA cannot be recovered. However, annotations are essential for many genomic analyses. We therefore employed a homology-based approach implemented in the program liftoff v1.6.1 to predict thylacine orthologs of Tasmanian devil genes (Shumate and Salzberg 2021). Exons from the Tasmanian devil RefSeq annotation were mapped to the thylacine genome assembly with minimap2 (Li 2018; O’Leary, et al. 2016). Thylacine gene models were then produced by linking mapped exons of a common parent feature, retaining only those which preserved the structure of their corresponding Tasmanian devil reference annotation (allowing a distance factor of 4X; parameter -d 4, Supplementary Table 6).

### Assembly Evaluation and Comparisons

Assembly completeness and integrity were assessed using Benchmarking Universal Single-Copy Orthologs annotated by BUSCO (v5.2.2) with the mammalian_odb10 ortholog database. These results were compared with several recent *de novo* marsupial genome assemblies (Fig. 1e, Supplementary Tables 3 and 4) (Brandies, et al. 2020; Dudchenko, et al. 2017; Johnson, et al. 2018; Peel, et al. 2022; Seppey, et al. 2019; Tian, et al. 2022). Comparison genomes were chosen to represent a variety of marsupial lineages and assembly approaches released within the past 4 years. Genome assembly metrics (Fig. 1d, Supplementary Table 2) were calculated using the stats.sh script in the BBmap package (v37.93) (Bushnell 2014).

### Heterozygosity

To calculate heterozygosity across species, short reads were aligned to each genome assembly with bwa-mem2 (-M flag; Supplementary Table 4) (Vasimuddin, et al. 2019). Samtools v1.11 was used to filter alignments (view -F 3340 -f 3) and remove duplicates (fixmate -m, markdup -r -S) (Li, et al. 2009). Pileups and variant filtering were performed using bcftools v1.11 mpileup (-q 20 -Q 20 -C 50) call (-m) and view (QUAL > 20, && DP>N && DP<M, where N and M represented 0.5x and 2x the average alignment coverage post-filtering) (Danecek, et al. 2021). Variants within repeats were identified with Red v2.0 and excluded using bedtools v2.27.1, due to low accuracy of read mapping within such regions (Girgis 2015; Quinlan and Hall 2010). This approach was applied to all genomes for this analysis rather than RepeatMasker alone, as Red has similar masking sensitivity to RepeatMasker with orders-of-magnitude lower computational overhead (Girgis 2015). Per-base heterozygosity was taken as the quotient of heterozygous positions and total callable genomic positions (Fig. 2b).

## Acknowledgements

We thank the Tasmanian Museum and Art Gallery and Museums Victoria for the use of the images in Fig. 1b & c respectively. This work is supported by Discovery Projects DP210102645 and DP210100505 from the Australian Research Council to AJP and SRF. CYF is supported on Ruth L. Kirschstein National Research Service Award 1F32GM139240-01 by the National Institute of General Medicinal Sciences of the National Institutes of Health. CYF performed all genomic analyses. AJP and SRF collected and sequenced fat-tailed dunnart samples used in heterozygosity analyses. CYF wrote the manuscript with input from all authors. We thank Elise Ireland for proofreading.

## Data Availability

Thylacine assembly, reads and inferred transcripts are submitted under NCBI BioProject PRJNA354646. Dunnart reads have been submitted to NCBI under SUB11101552.

